# SLC4A11 mediates ammonia import and promotes cancer stemness in hepatocellular carcinoma

**DOI:** 10.1101/2024.08.06.606899

**Authors:** Ameer L. Elaimy, Marwa O. El-Derany, Jadyn James, Zhuwen Wang, Ashley N. Pearson, Erin A. Holcomb, Amanda K. Huber, Miguel Gijón, Hannah N. Bell, Viraj R. Sanghvi, Timothy L. Frankel, Grace L. Su, Elliot B. Tapper, Andrew W. Tai, Nithya Ramnath, Christopher P. Centonze, Irina Dobrosotskaya, Julie A. Moeller, Alex K. Bryant, David A. Elliott, Enid Choi, Joseph R. Evans, Kyle C. Cuneo, Thomas J. Fitzgerald, Daniel R. Wahl, Meredith A. Morgan, Daniel T. Chang, Max S. Wicha, Theodore S. Lawrence, Yatrik M. Shah, Michael D. Green

## Abstract

End stage liver disease is marked by portal hypertension, systemic elevations in ammonia, and development of hepatocellular carcinoma (HCC). While these clinical consequences of cirrhosis are well described, it remains poorly understood whether hepatic insufficiency and the accompanying elevations in ammonia contribute to HCC carcinogenesis. Using preclinical models, we discovered that ammonia entered the cell through the transporter SLC4A11 and served as a nitrogen source for amino acid and nucleotide biosynthesis. Elevated ammonia promoted cancer stem cell properties *in vitro* and tumor initiation *in vivo*. Enhancing ammonia clearance reduced HCC stemness and tumor growth. In patients, elevations in serum ammonia were associated with an increased incidence of HCC. Taken together, this study forms the foundation for clinical investigations using ammonia lowering agents as potential therapies to mitigate HCC incidence and aggressiveness.

## Introduction

The vast majority of hepatocellular carcinoma (HCC) cases arise in the setting of chronic liver disease and cirrhosis (1). Given the high risk of developing HCC in patients with cirrhosis, screening programs are recommended to detect tumors at an early stage and guide clinical decision-making (2). Most patients with liver-confined HCC are not candidates for potentially curative resection or transplant due to poor functional status, inadequate hepatic reserve, or tumor location (3–6). These patients are subsequently evaluated for various liver-directed therapies (7). Although local treatments are generally effective in providing tumor control, the development of new intrahepatic lesions is common (8). Cell damage from oxidative stress caused by chronic inflammation is thought to contribute to the high propensity to initiate tumors in cirrhotic patients (9–11). However, a greater understanding of the impact of the liver microenvironment on HCC initiation is needed with the goal of potentially exploiting the pathophysiological consequences that result from liver dysfunction for therapeutic benefit.

Ammonia is a nitrogenous waste product of amino acid metabolism that undergoes detoxification by the liver via the urea cycle (12). In patients with cirrhosis, inadequate ammonia clearance contributes to neuropsychiatric sequelae including hepatic encephalopathy (13). While well recognized as a toxic metabolic byproduct, some malignancies can metabolically recycle ammonia for incorporation into anabolic pathways that promote tumor growth and contribute to therapy resistance (14, 15). These observations are of particular relevance to HCC as it arises in a background of cirrhosis (16). However, whether ammonia contributes to HCC initiation and progression remains unknown.

Cancer stem cells (CSCs) are a subpopulation of cells that are responsible for tumor initiation, therapy resistance, and disease recurrence (17, 18). CSCs reside in a niche of immune and stromal cells, extracellular matrix contacts, and secretory molecules that, together, support self-renewal (19). HCC is enriched in CSCs that are adapted to survive in the unique microenvironment of a cirrhotic liver (20–23). In this study, we explored the interplay between metabolic alterations in cirrhosis and hepatocellular carcinogenesis using preclinical models and patient data. We discovered that ammonia contributed to tumor initiation by serving as a precursor for amino acid and nucleotide biosynthesis in CSCs, and that elevated ammonia correlated with cancer incidence and poor prognosis in HCC patients. Our work suggests that targeting this crosstalk may reduce the incidence and aggressiveness of HCC in patients with chronic liver disease.

## Results

### Elevated ammonia is correlated with an increased incidence of hepatocellular carcinoma in cirrhotic patients

We and others have shown that ammonia can support tumor growth in breast and colorectal cancer (14, 15). It remains unclear whether hepatic insufficiency and longitudinal elevations in ammonia contribute to hepatocellular carcinogenesis. To examine this question, we identified a cohort of 363,032 Veterans with cirrhosis diagnosed between the years 1999 and 2024. Approximately 48,476 patients within this cohort had quantification of serum ammonia levels at the time of cirrhosis diagnosis (Figure 1A), and 9,441 of these patients later developed HCC with a median time to diagnosis of 3.1 years (Cohort 1). We first asked whether elevations in baseline ammonia were associated with an increased risk of developing HCC. Interestingly, we observed a significant positive association between baseline ammonia levels and HCC incidence (Figure 1B). To evaluate this association more rigorously, we next stratified patients by baseline ammonia level into those patients with high and low mean ammonia values. Cirrhotic patients with high and low ammonia were similar in terms of age, ethnicity, and etiology of their underlying liver disease (Table 1). We observed that patients with high baseline ammonia had a significantly higher cumulative incidence of HCC (HR 2.02 [95% CI, 1.86-2.20], *p* < 0.0001) as compared to patients with low ammonia levels (Figure 1C).

**Figure 1.**
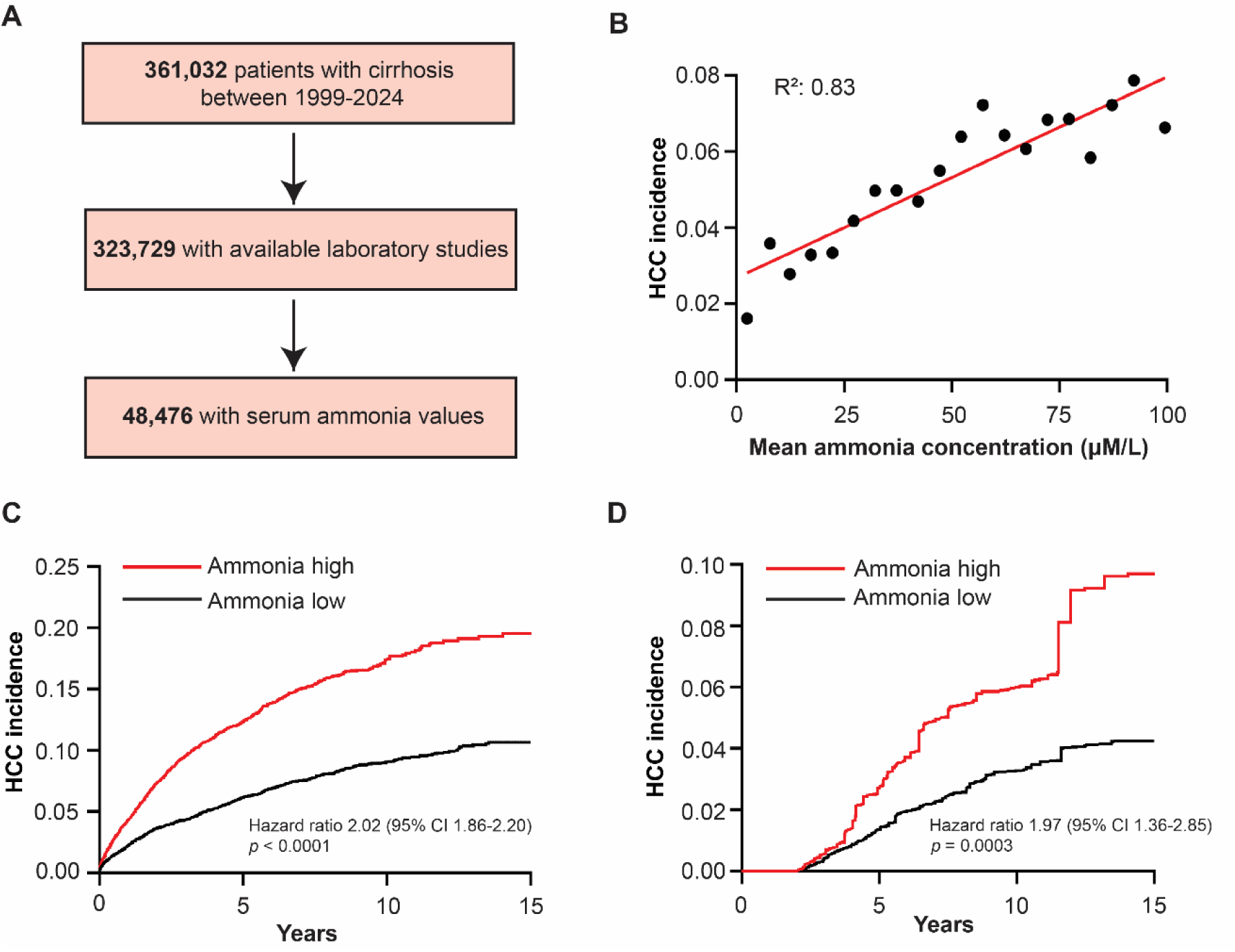
Elevated ammonia is correlated with an increased incidence of hepatocellular carcinoma in cirrhotic patients. (**A**) CONSORT diagram of patient population. (**B**) Scatter plot depicting correlation between mean ammonia concentration and HCC incidence fit using linear regression. (**C**) Cumulative HCC incidence stratified by mean ammonia levels (ammonia high, *n =* 20,099; ammonia low, *n* = 28,377). (**D**) Two-year landmark analysis of HCC incidence in patients with high and low ammonia using propensity score matching. Hazard ratio log-rank test, *p* values, and 95% confidence intervals indicated.

**Table 1:** Patient population baseline characteristics of cirrhotic cohort.

| Variable | Total<br>(48,476)<br>No. | % | Ammonia low<br>(28,377)<br>No. | % | Ammonia high<br>(20,099)<br>No. | % |
| --- | --- | --- | --- | --- | --- | --- |
| **Age, years** |  |  |  |  |  |  |
| Median (IQR) | 61.1 (8.4) |  | 61.6 (8.2) |  | 60.3 (8.6) |  |
| **Male sex** | 46,726 | 96.4 | 27,305 | 96.2 | 19,421 | 96.6 |
| **Ethnicity** |  |  |  |  |  |  |
| White | 30,634 | 63.2 | 18,082 | 63.7 | 12,552 | 62.5 |
| Black | 9,173 | 18.9 | 5,618 | 19.8 | 3,555 | 17.7 |
| Hispanic | 3,794 | 7.8 | 1,900 | 6.7 | 1,894 | 9.4 |
| Other/unknown | 4,875 | 10.1 | 2,777 | 9.8 | 2,098 | 10.4 |
| **Underlying liver disease** |  |  |  |  |  |  |
| Viral | 9,859 | 20.3 | 5,517 | 19.4 | 4,342 | 21.6 |
| Alcohol | 28,437 | 58.7 | 16,690 | 58.8 | 11,747 | 58.4 |
| MASLD | 5,822 | 12.0 | 3,392 | 12.0 | 2,430 | 12.1 |
| Other | 4,358 | 9.0 | 2,778 | 9.8 | 1,580 | 7.9 |
| **ALBI Grade** |  |  |  |  |  |  |
| 1 | 9,299 | 19.2 | 7,361 | 25.9 | 1,938 | 9.6 |
| 2 | 24,508 | 50.6 | 14,600 | 51.5 | 9,908 | 49.3 |
| 3 | 14,669 | 30.3 | 6,416 | 22.6 | 8,253 | 41.1 |
| **MELD Score** |  |  |  |  |  |  |
| ≤ 9 | 13,484 | 27.8 | 9,309 | 32.8 | 4,175 | 20.8 |
| 10-19 | 23,809 | 49.1 | 13,448 | 47.4 | 10,361 | 51.5 |
| ≥ 20 | 11,183 | 23.1 | 5,620 | 19.8 | 5,563 | 27.7 |
| **eCTP Class** |  |  |  |  |  |  |
| A | 18,848 | 38.9 | 13,964 | 49.2 | 4,884 | 24.3 |
| B | 18,571 | 38.3 | 10,132 | 35.7 | 8,439 | 42.0 |
| C | 11,057 | 22.8 | 4,281 | 15.1 | 6,776 | 33.7 |

Abbreviations: ALBI, albumin-bilirubin; eCTP, electronic Child-Turcotte-Pugh; IQR, interquartile range; MASLD, metabolic dysfunction-associated steatotic disease; MELD, model for end-stage liver disease

To understand whether elevations in ammonia correlated with increased cumulative incidence of HCC in all subsets of patients, we performed additional analyses. Albumin-bilirubin (ALBI) grade (24), Model for End-Stage Liver Disease (MELD) (25), and Child-Turcotte-Pugh class (CTP) (26) are widely utilized assessments of the degree of liver dysfunction in patients with cirrhosis. We observed that elevations in baseline ammonia correlated with increased HCC incidence regardless of the ALBI grade, MELD score, and electronic CTP (eCTP) class (Supplementary Figure 1A). To more rigorously evaluate this association, we next conducted a propensity score weighted landmarked analysis. Multivariable analysis showed that the cumulative incidence of HCC was similar in patients with cirrhosis who had ammonia quantified as compared to those patients who did not have ammonia quantified (Supplemental Figure 1B). In contrast, multivariable analysis showed that the cumulative incidence of HCC remained significantly higher in patients with elevated ammonia as compared to those patients without elevations after correction for clinicopathologic features (HR 1.97 [95% CI, 1.36-2.85], *p* = 0.0003) (Figure 1D). Together, these data suggest that elevations in serum ammonia in patients with cirrhosis may be associated with a higher risk of developing HCC.

### Ammonia contributes to cancer stem cell function and hepatocellular carcinoma initiation

Given our observation that elevated ammonia is associated with a higher HCC incidence, we hypothesized that ammonia may regulate CSCs because they are responsible for tumor initiation and rely on a unique microenvironment to sustain their function (20). Three-dimensional sphere culture enriches a population of cells with self-renewal properties and is one method to quantify CSCs *in vitro* (27). To initially test this hypothesis, we evaluated whether ammonia influenced establishment of hepatospheres. We treated the human HCC cell line HepG2 with 10 mM ammonium chloride because that is within the physiological concentration of ammonia in the liver, the concentration we observed peak hepatosphere formation in HepG2 cells (Supplementary Figure 2A), and has been used in prior studies studying ammonia in cancer (16). We observed that ammonium chloride treatment increased hepatosphere number and diameter in HepG2 cells (Figure 2A). To extend this finding, we next evaluated a second human HCC cell line, HUH7. Again, ammonium chloride treatment increased hepatosphere number and diameter (Figure 2B). Finally, we utilized a murine HCC cell line derived from tumors generated by overexpression of Myc and knockout of p53 as previously described (hereafter termed mHCC) (28). Indeed, we observed that ammonium chloride treatment increased hepatosphere number and diameter in mHCC cells (Figure 2C). These data provide evidence that ammonia promotes hepatosphere formation and growth.

**Figure 2.**
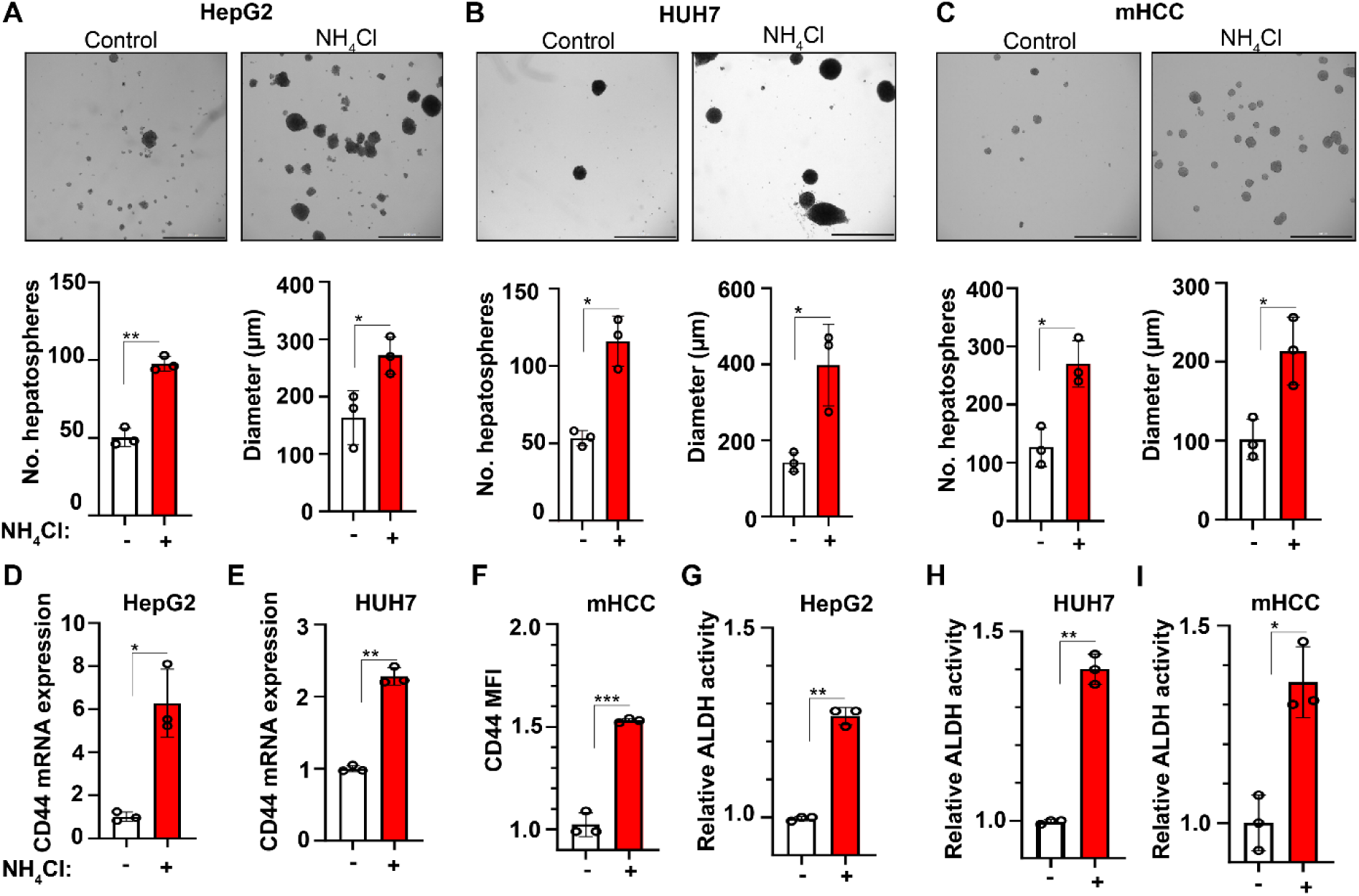
Ammonia promotes the acquisition of cancer stem cell properties *in vitro.* Representative brightfield micrographs, number of hepatospheres, and diameter of hepatospheres formed with and without ammonium chloride (10 mM) in (**A**) HepG2, (**B**) HUH7, and (**C**) mHCC cells. Mean ± SD (*n* = 7-9). Scale bar represents 1000 µm. CD44 mRNA expression in (**D**) HepG2 and (**E**) HUH7 cells with and without ammonium chloride (10 mM). Mean ± SD (*n* = 3). (**F**) CD44 surface expression by flow cytometry in control and ammonium chloride (10 mM) treated mHCC hepatospheres. Mean fluorescence intensity (MFI) fold change ± SD (*n* = 3). ALDH activity in (**G**) HepG2, (**H**) HUH7, and (**I**) mHCC hepatospheres with and without ammonium chloride (10 mM) (*n* = 3). * *p* ≤ 0.05, ** *p* ≤ 0.005, *** *p* ≤ 0.0005 by two-tailed *t* test.

CD44 is a well-established marker of CSCs in HCC and other systems (29–32). Ammonium chloride treatment increased CD44 mRNA expression in HepG2 (Figure 2D) and HUH7 hepatospheres (Figure 2E). Ammonium chloride treatment also increased CD44 surface expression in mHCC hepatospheres (Figure 2F and Supplementary Figure 2B). The enzymatic activity of aldehyde dehydrogenase (ALDH) is elevated in HCC CSCs and is another method for quantifying stemness *in vitro* (33, 34). Ammonium chloride treatment increased ALDH activity in HepG2 (Figure 2G), HUH7 (Figure 2H), and mHCC (Figure 2I) hepatospheres. These data suggest that ammonia promotes cancer stemness in HCC *in vitro*.

Limiting dilution analysis is the gold-standard method for quantifying tumor initiation *in vivo* (35, 36). We therefore treated HepG2 hepatospheres with ammonium chloride and conducted limiting dilution tumor initiation studies in NOD *scid* gamma (NSG) mice. We observed that ammonium chloride treatment increased the frequency of tumor initiating cells (TICs) (Figure 3A). We next repeated limiting dilution analysis with mHCC cells. Again, ammonium chloride treatment increased the frequency of TICs (Figure 3B). We also observed that *ex-vivo* ammonium chloride treatment of HepG2 hepatospheres prior to inoculation in NSG mice increased tumor volume (Figure 3C and Supplementary Figure 3A) and tumor weight (Figure 3D and Supplementary Figure 3B). Similarly, *ex-vivo* ammonium chloride treatment of mHCC hepatospheres prior to inoculation in NSG mice increased tumor volume (Figure 3E and Supplementary Figure 3C) and tumor weight (Figure 3F and Supplementary Figure 3D). These data indicate that ammonia increases tumor initiation and growth in HCC *in vivo*.

**Figure 3.**
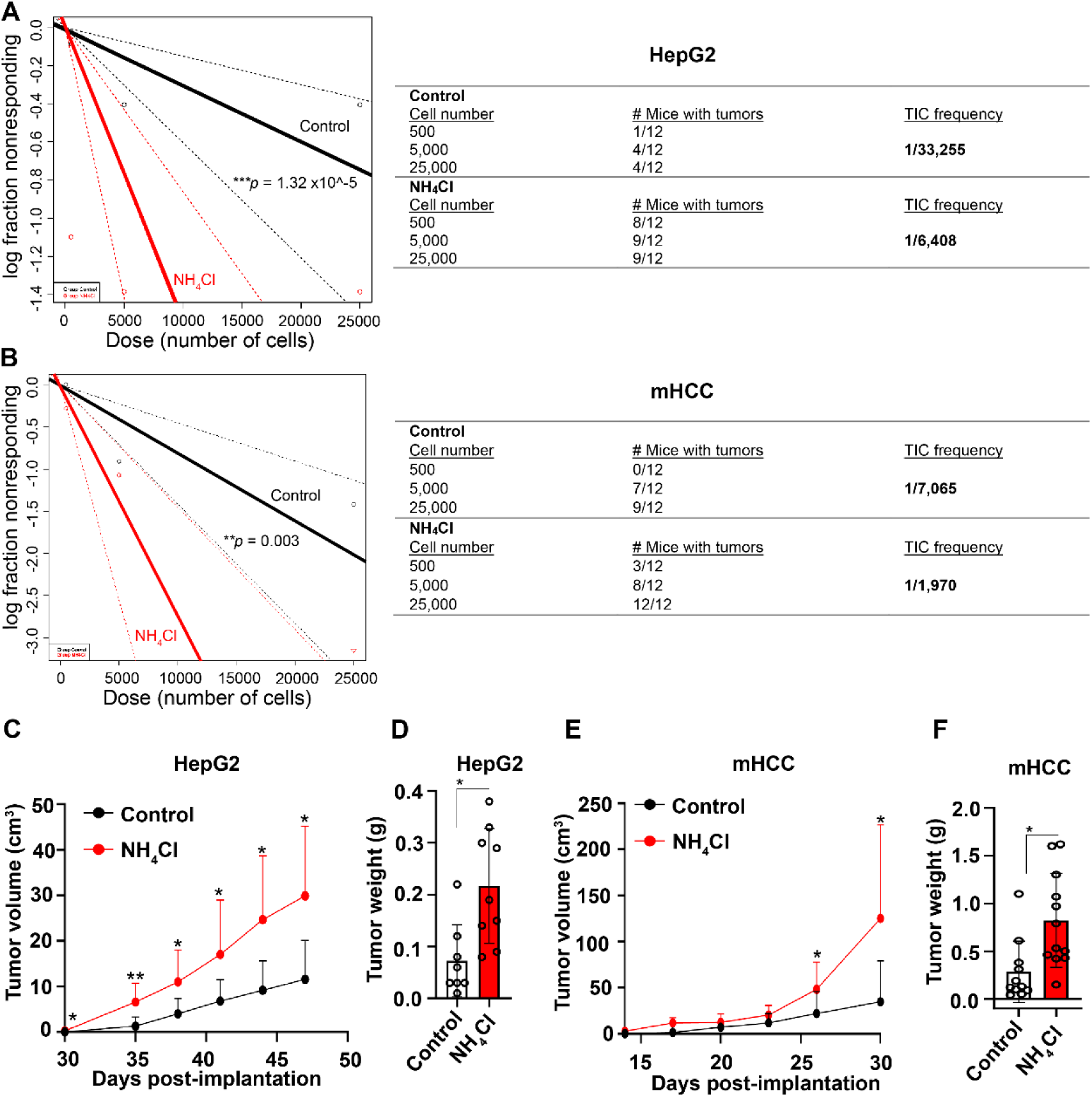
Ammonia contributes to tumor initiation *in vivo*. Control and ammonium chloride (10 mM) treated hepatospheres derived from (**A**) HepG2 and (**B**) mHCC cells were dissociated into single cells and implanted into NSG mice at the indicated cell numbers (*n* = 12 tumors per arm), and tumor initiation and TIC frequency was quantified using extreme limiting dilution analysis. HepG2 (**C**) tumor volume and (**D**) tumor weight was quantified in the 500-cell titration group. mHCC (**E**) tumor volume and (**F**) tumor weight was quantified in the 500-cell titration group. Data are represented as means ± SD. * *p* ≤ 0.05, ** *p* ≤ 0.005 by two-tailed *t* test.

### SLC4A11 is upregulated in hepatocellular carcinoma stem cells and functions as an ammonia importer

Next, we aimed to determine the mechanism by which ammonia promotes stemness and tumor initiation. A recent report demonstrated that the ammonia transporter SLC4A11 promotes HCC growth and is associated with poor prognosis in patients (37). We therefore hypothesized that the transport of ammonia may have important functional consequences on influencing the behavior of CSCs. Indeed, SLC4A11 mRNA expression was upregulated in HepG2 and HUH7 hepatospheres when compared to adherent cells (Figure 4A) and further induced following ammonium chloride treatment in HepG2 and HUH7 hepatospheres (Figure 4B).

**Figure 4.**
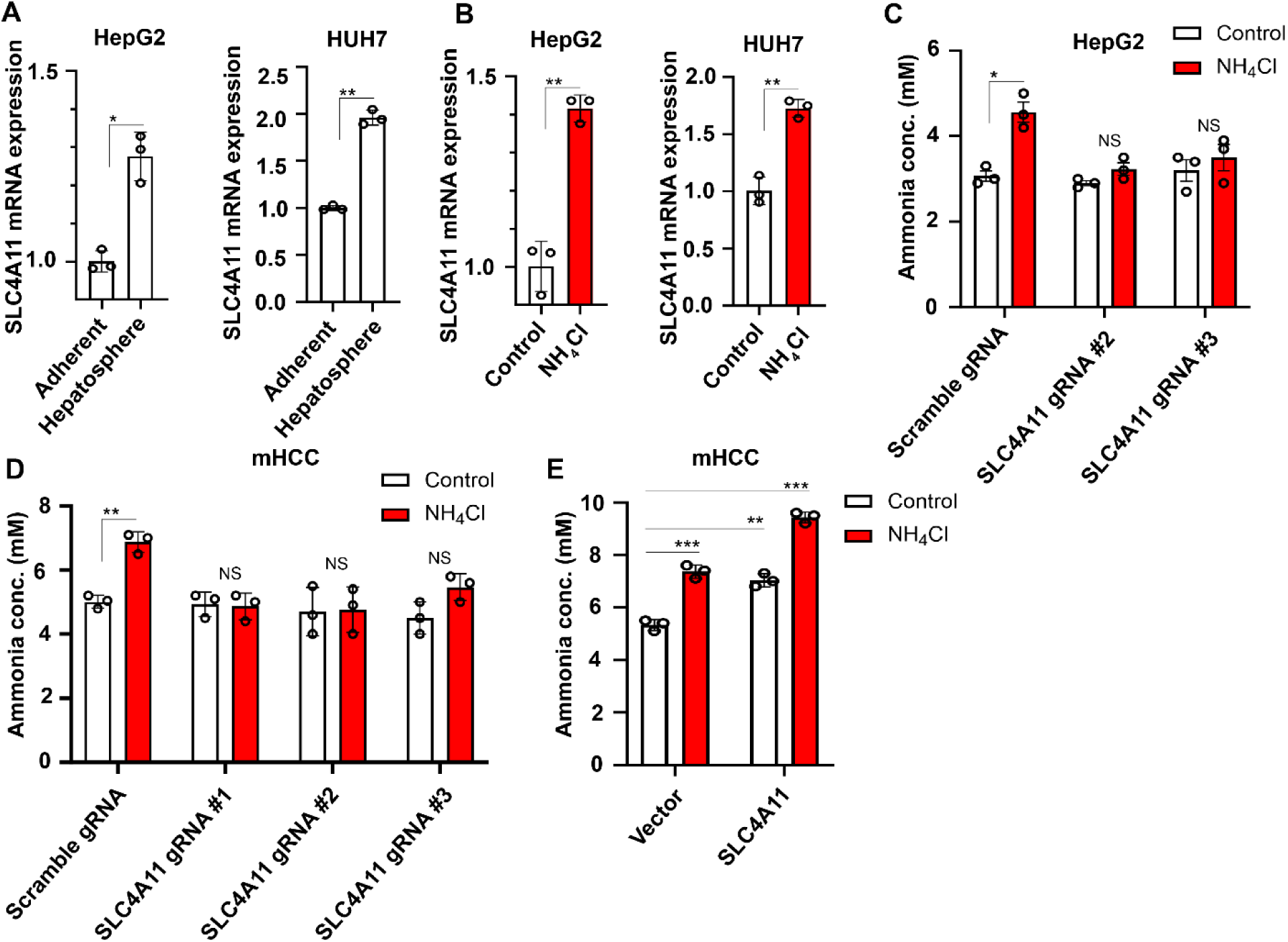
SLC4A11 functions as an ammonia importer in hepatocellular carcinoma stem cells. (**A**) SLC4A11 mRNA expression was quantified in 2D vs 3D culture of HepG2 (left) and HUH7 (right) cells. Mean ± SD (*n* = 3). (**B**) SLC4A11 mRNA expression was quantified in HepG2 (left) and HUH7 (right) hepatospheres with and without ammonium chloride (10 mM). Mean ± SD (*n* = 3). SLC4A11 was depleted in (**C**) HepG2 and (**D**) mHCC cells by Crispr/Cas9 using 2-3 independent gRNAs and ammonia concentration in control and SLC4A11 KO hepatospheres with and without ammonium chloride (10 mM) was quantified. Mean ± SD (*n* = 3). (**E**) tdTomato-tagged SLC4A11 was ectopically expressed in mHCC cells and ammonia concentration in control and SLC4A11 overexpression hepatospheres with and without ammonium chloride (10 mM) was quantified. Mean ± SD (*n* = 3). *p* ≤ 0.05, ** *p* ≤ 0.005, *** *p* ≤ 0.0005 by two-tailed *t* test.

SLC4A11 is known to transport ammonia both intra- and extracellularly (37–39). To determine the directionality of ammonia transport in HCC CSCs, we used Crispr/Cas9 to knock out (KO) SLC4A11 in HepG2 and mHCC cells. We designed 3 candidate human and murine gRNAs and proceeded with experiments with gRNAs that generated complete KO of SLC4A11 (human gRNAs #2 and 3 and murine gRNAs #1-3) (Supplementary Figure 4, A-D). We observed an increase in intracellular ammonia concentration in control HepG2 and mHCC hepatospheres treated with ammonium chloride, but did not observe an increase in SLC4A11 KO cells (Figure 4, C and D). Moreover, ectopic expression of SLC4A11 resulted in an increase in intracellular ammonia concentration in mHCC hepatospheres (Figure 4E and Supplementary Figure 4E). These data suggest that SLC4A11 functions as an ammonia importer in HCC CSCs.

To further assess the potential contribution of SLC4A11-mediated ammonia import in inducing stemness, we quantified hepatosphere number in control and SLC4A11 KO mHCC hepatospheres. Ammonium chloride treatment increased hepatosphere number in control cells, but it did not in SLC4A11 KO cells (Figure 5A). Similar results were obtained in HepG2 cells (Figure 5B). Furthermore, ectopic expression of SLC4A11 in mHCC cells resulted in an increase in hepatosphere number (Figure 5C). To provide additional evidence of a causal role of SLC4A11 in promoting stemness, we observed a reduction in CD44 mRNA expression (Figure 5D) and ALDH activity (Figure 5E) in SLC4A11 KO mHCC hepatospheres in the presence of ammonium chloride.

**Figure 5.**
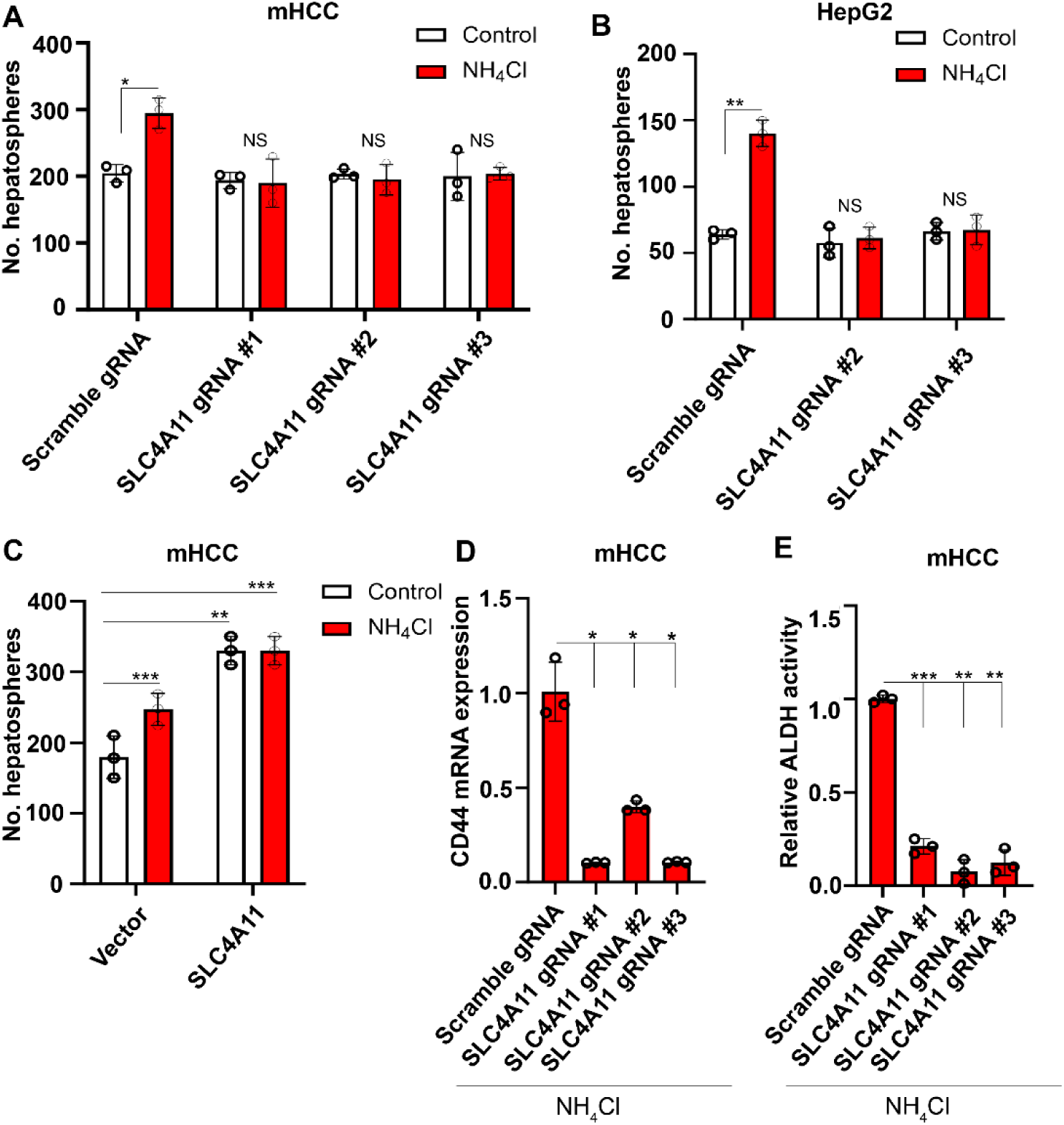
SLC4A11-mediated ammonia transport sustains a cancer stem cell phenotype. Hepatosphere number in control and SLC4A11 KO (**A**) mHCC and (**B**) HepG2 cells with and without ammonium chloride (10 mM). Mean ± SD (*n* = 3). (**C**) Hepatosphere number in control and SLC4A11 overexpressing mHCC cells with and without ammonium chloride (10 mM). Mean ± SD (*n* = 3). (**D**) CD44 mRNA expression and (**E**) ALDH activity in control and SLC4A11 KO mHCC hepatospheres in the presence of ammonium chloride (10 mM). Mean ± SD (*n* = 3). * *p* ≤ 0.05, ** *p* ≤ 0.005, *** *p* ≤ 0.0005 by two-tailed *t* test.

### Ammonia augments amino acid and nucleotide biosynthesis in a SLC4A11 dependent manner

An important question arising from our data is the fate of ammonia-derived nitrogen in HCC CSCs. Since we established that SLC4A11 mediates intracellular ammonia transport in CSCs, we hypothesized that ammonia incorporates into metabolic pathways that contribute to tumor growth. We therefore performed unbiased tracing of the nitrogen metabolome using high-performance liquid chromatography-mass spectrometry (HPLC-MS) and assessed the fate of ^15^NH_4_Cl in control and SLC4A11 KO HepG2 hepatospheres by characterizing a panel of 215 isotopologues of nitrogen as previously described (Figure 6A) (15). This analysis found that ammonia-derived nitrogen primarily enters central biosynthetic pathways of amino acids and nucleotides (Figure 6B, Supplementary Figure 5A, and Supplementary Table 1). Using metabolite set enrichment analysis (MSEA), we identified 15 pathways by the Kyoto Encyclopedia of Genes and Genomes (KEGG) and 14 pathways by the Small Molecule Pathway Database (SMPDB) that were significantly enriched with ^15^NH_4_Cl treatment (Figure 6C and Supplementary Figure 5B). Isotopologues of the amino acids aspartate, glutamine, alanine, and glutamate and the nucleotides guanine, guanosine, uridine, uracil, and adenine were of the most enriched metabolites (Figure 6, D and E). These data suggest that ammonia-derived nitrogen is incorporated into glutamine that serves as an intermediary for the synthesis of other amino acids and nucleotides. Importantly, ^15^N labeling in SLC4A11 KO HepG2 hepatospheres was significantly reduced compared to controls further substantiating our conclusion that SLC4A11 serves as a critical intracellular ammonia transporter in HCC CSCs (Figure 6, B, D, and E).

**Figure 6.**
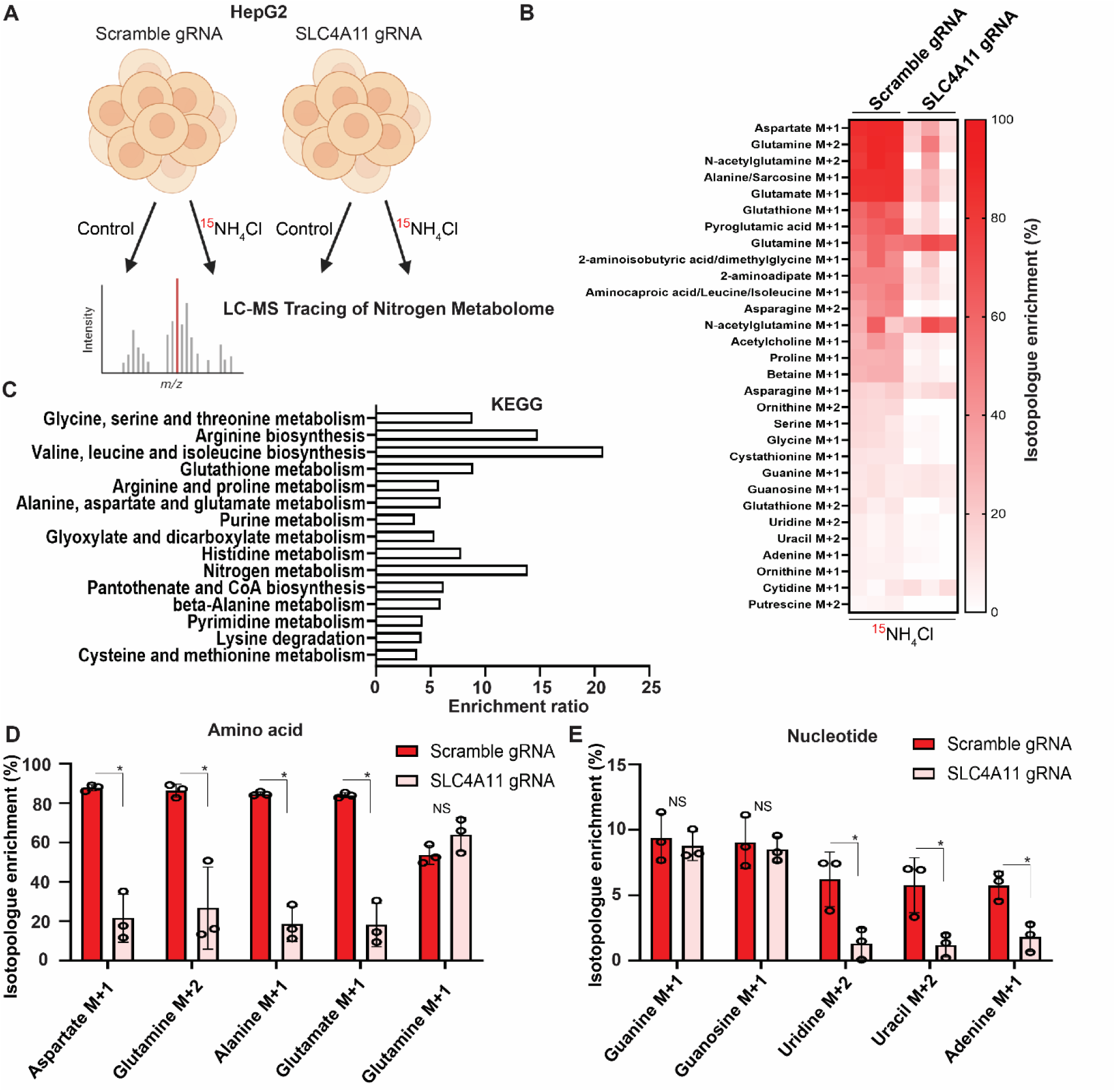
Ammonia augments amino acid and nucleotide biosynthesis in a SLC4A11 dependent manner. (**A**) Schematic depicting LC-MS based nitrogen tracing in control and SLC4A11 KO HepG2 hepatospheres with and without 8 hours of ammonium chloride (10 mM) (created with BioRender.com). (**B**) Isotopologue enrichment of top 30 metabolites in control and SLC4A11 KO HepG2 hepatospheres (background subtracted from ^15^NH_4_Cl treated samples, *n* = 3). (**C**) KEGG gene set enrichment analysis for 15 pathways significantly enriched by hypergeometric testing (*n* = 3). Normalized isotopologue enrichment of representative (**D**) amino acids and (**E**) nucleotides in control and SLC4A11 KO HepG2 hepatospheres. * *p* ≤ 0.05, ** *p* ≤ 0.005, *** *p* ≤ 0.0005 by two-tailed *t* test.

### Ammonia clearance reduces tumorigenesis in vivo

We established that ammonia promotes tumor initiation and growth in HCC by impacting the function of CSCs. Using a spontaneous HCC model generated by hydrodynamic transfection of Myc, gp53/Cas9, and sleeping beauty transposase for stable genomic integration in C57BL/6J mice as previously described (28), we observed that murine livers with HCC have elevated ammonia concentrations in tumors when compared to normal control livers (Figure 7A). Ornithine is an amino acid that augments urea cycle flux to promote ammonia clearance, is used clinically to treat cirrhotic patients (40), and we have previously shown that ornithine can lower intratumoral ammonia within the liver (14). We therefore tested the effects of ammonia clearance using ornithine in mice with HCCs generated by hydrodynamic transfection and observed a striking reduction in tumor growth as measured by liver weight (Figure 7B). Ornithine treatment at physiologically relevant doses also reduced intratumoral ammonia concentration (Figure 7C), CD44 mRNA expression (Figure 7D), SLC4A11 mRNA expression (Figure 7E), and ALDH activity (Figure 7F). To further test the effects of ammonia on tumor growth, we established mHCC xenografts and found that mice fed a high ammonia diet exhibited increased tumor weights (Figure 7G) and intratumoral ammonia concentrations compared to control mice (Figure 7H). Importantly, ornithine treatment in mice fed a high ammonia diet resulted in smaller tumor weights and lower intratumoral ammonia concentrations. These experiments demonstrate that ammonia clearance reduces CSC properties and tumor growth *in vivo*.

**Figure 7.**
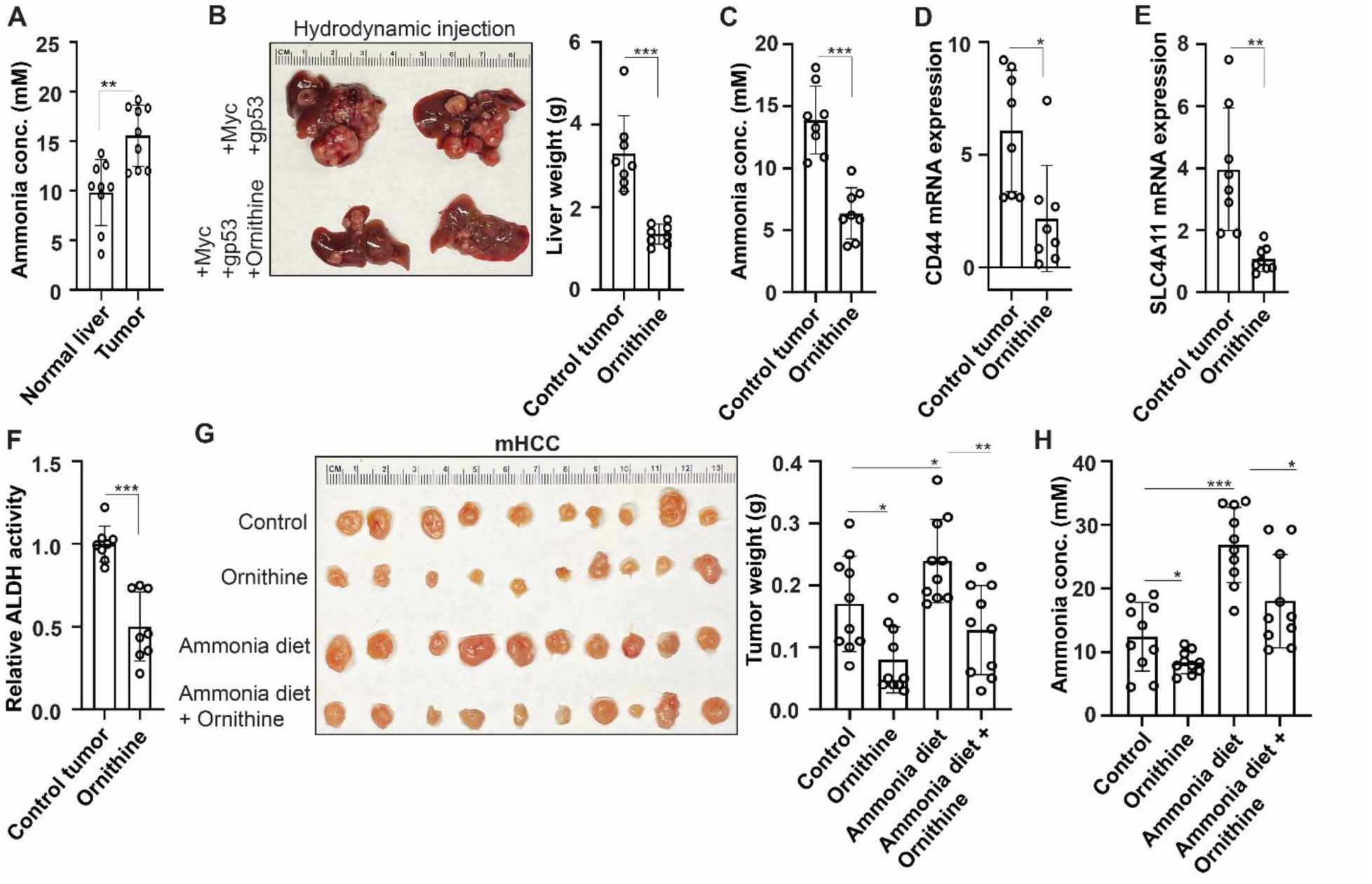
Ammonia clearance reduces tumor burden and cancer stem cell markers *in vivo*. (**A**) Ammonia concentration in livers of control and HCC tumor-bearing C57BL/6J mice following hydrodynamic transfection of Myc, gp53/Cas9, and sleeping beauty transposase. Mean ± SD (*n* = 9 per arm). (**B**) Representative photo and liver weight, (**C**) ammonia concentration, (**D**) CD44 mRNA expression, (**E**) SLC4A11 mRNA expression, and (**F**) ALDH activity of tumor bearing C57BL/6J mice generated by hydrodynamic tail vein injection with and without ornithine treatment. Mean ± SD (*n* = 8 per arm). **(G)** Representative photo and tumor weight, and **(H)** ammonia concentration of mHCC tumors established in NSG mice with the indicated conditions. Mean ± SD (*n* = 10 per arm). * *p* ≤ 0.05, ** *p* ≤ 0.005, *** *p* ≤ 0.0005 by two-tailed *t* test.

### Elevated ammonia is associated with poor prognosis in hepatocellular carcinoma patients

Given our finding that ammonia promotes tumor initiation and growth in preclinical models, we hypothesized that elevations in ammonia may be associated with adverse outcomes in HCC patients. To evaluate this, we identified a cohort of 3,550 patients with a diagnosis of HCC from the cancer registry and similarly stratified patients with high and low ammonia (Cohort 2). HCC patients with high and low ammonia were similar in terms of age, ethnicity, and etiology of their underlying liver disease, and patients with high ammonia had worse overall liver function as measured by ALBI grade, MELD score, and eCTP class (Table 2). On univariate analysis, we observed that patients with elevated ammonia had a worse overall survival (OS) (HR 1.29 [95% CI, 1.20-1.38], *p* < 0.0001) (Figure 8A). On subset analysis, patients with elevated ammonia had significantly inferior OS in the subset of patients with relatively well-preserved liver function (ALBI grade 1, MELD ≤ 9, and eCTP class A) (Figure 8B). There was a trend towards inferior OS in those patients with moderate liver function and elevated ammonia (ALBI grade 2, MELD score 10-19). Multivariable modeling using propensity score weighting confirmed that patients with elevated ammonia had a worse OS (HR 1.26 [95% CI, 1.14-1.38], *p* < 0.0001) (Figure 8C). These data suggest that elevated ammonia is associated with poor prognosis in patients with HCC.

**Figure 8.**
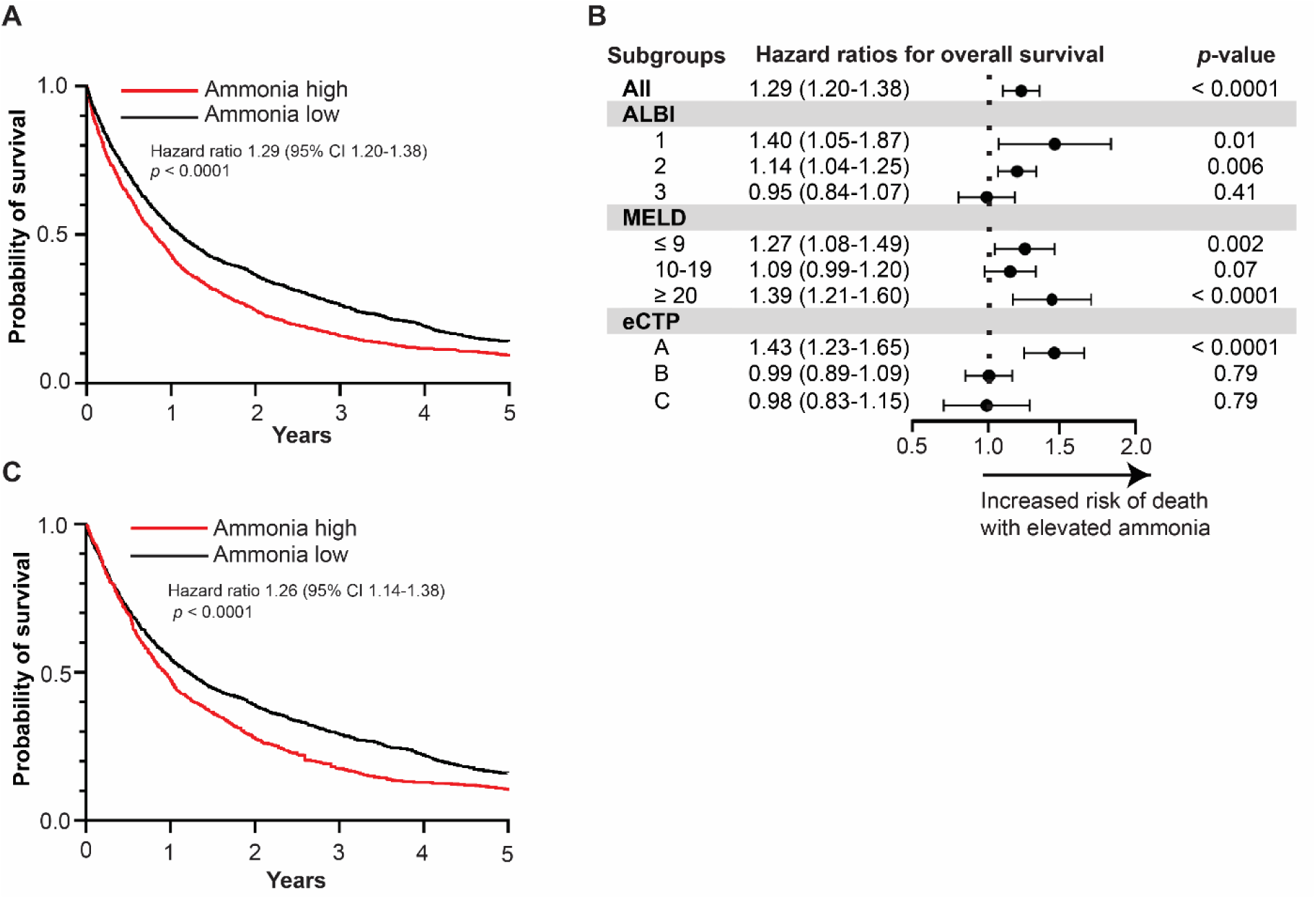
Elevated ammonia is associated with poor overall survival in hepatocellular carcinoma patients. (**A**) OS of patients with HCC stratified by mean ammonia levels (ammonia high, *n =* 1,813; ammonia low, *n* = 1,737). (**B**) OS of patients diagnosed with HCC by ALBI grade, MELD score, and eCTP class. (**C**) OS of patients with high and low ammonia using propensity score matching. Hazard ratio log-rank test, *p* values, and 95% confidence intervals indicated.

**Table 2:**
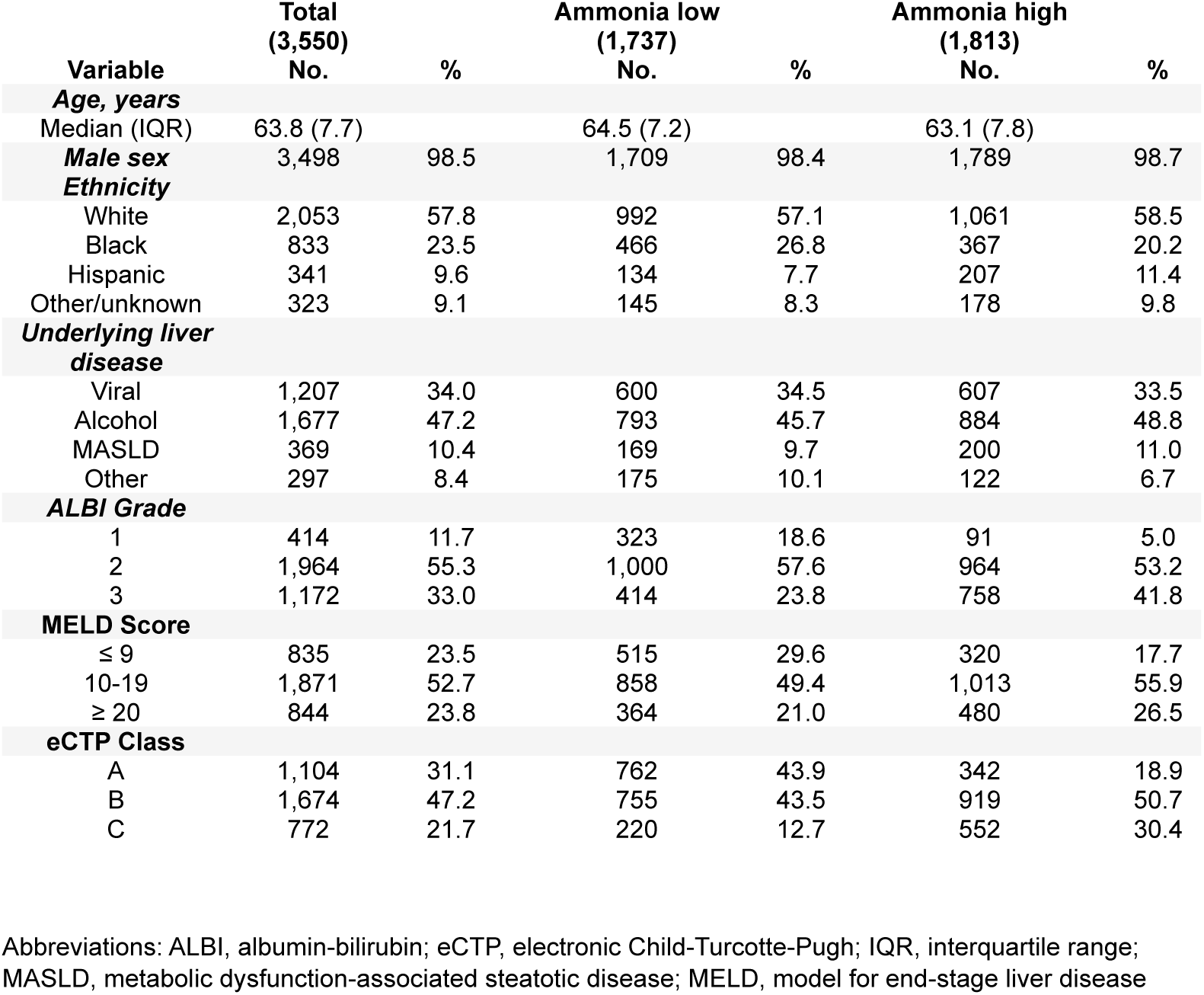
Patient population baseline characteristics of HCC cohort.

| Variable | Total<br>(3,550)<br>No. | % | Ammonia low<br>(1,737)<br>No. | % | Ammonia high<br>(1,813)<br>No. | % |
| --- | --- | --- | --- | --- | --- | --- |
| **Age, years** |  |  |  |  |  |  |
| Median (IQR) | 63.8 (7.7) |  | 64.5 (7.2) |  | 63.1 (7.8) |  |
| **Male sex** | 3,498 | 98.5 | 1,709 | 98.4 | 1,789 | 98.7 |
| **Ethnicity** |  |  |  |  |  |  |
| White | 2,053 | 57.8 | 992 | 57.1 | 1,061 | 58.5 |
| Black | 833 | 23.5 | 466 | 26.8 | 367 | 20.2 |
| Hispanic | 341 | 9.6 | 134 | 7.7 | 207 | 11.4 |
| Other/unknown | 323 | 9.1 | 145 | 8.3 | 178 | 9.8 |
| **Underlying liver disease** |  |  |  |  |  |  |
| Viral | 1,207 | 34.0 | 600 | 34.5 | 607 | 33.5 |
| Alcohol | 1,677 | 47.2 | 793 | 45.7 | 884 | 48.8 |
| MASLD | 369 | 10.4 | 169 | 9.7 | 200 | 11.0 |
| Other | 297 | 8.4 | 175 | 10.1 | 122 | 6.7 |
| **ALBI Grade** |  |  |  |  |  |  |
| 1 | 414 | 11.7 | 323 | 18.6 | 91 | 5.0 |
| 2 | 1,964 | 55.3 | 1,000 | 57.6 | 964 | 53.2 |
| 3 | 1,172 | 33.0 | 414 | 23.8 | 758 | 41.8 |
| **MELD Score** |  |  |  |  |  |  |
| ≤ 9 | 835 | 23.5 | 515 | 29.6 | 320 | 17.7 |
| 10-19 | 1,871 | 52.7 | 858 | 49.4 | 1,013 | 55.9 |
| ≥ 20 | 844 | 23.8 | 364 | 21.0 | 480 | 26.5 |
| **eCTP Class** |  |  |  |  |  |  |
| A | 1,104 | 31.1 | 762 | 43.9 | 342 | 18.9 |
| B | 1,674 | 47.2 | 755 | 43.5 | 919 | 50.7 |
| C | 772 | 21.7 | 220 | 12.7 | 552 | 30.4 |

Abbreviations: ALBI, albumin-bilirubin; eCTP, electronic Child-Turcotte-Pugh; IQR, interquartile range; MASLD, metabolic dysfunction-associated steatotic disease; MELD, model for end-stage liver disease

## Discussion

The results of this study demonstrate that ammonia contributes to the function of CSCs and promotes tumor initiation in HCC. In preclinical models, the transporter SLC4A11 sustained high levels of intracellular ammonia and is at the nexus of a network that rewired the nitrogen metabolome to promote amino acid and nucleotide biosynthesis in HCC CSCs. In cirrhotic patients, elevated ammonia was associated with higher HCC incidence when correcting for liver function and using propensity score matching. Additionally, in patients with HCC, elevated ammonia was associated with worse overall survival. These findings increase our understanding of the role of ammonia in tumorigenesis. They also have substantial clinical implications because multiple pharmacologic agents that are commonly used to promote ammonia clearance in cirrhotic patients at ranges established to have clinical impact could be repurposed to be tested as therapeutic or preventative agents in trials.

A major finding of this study is that ammonia contributes to tumor initiation and growth in HCC by regulating CSC function. This observation is important as dysregulated ammonia metabolism is a common clinical finding in patients with cirrhosis, which is the most significant risk factor in developing HCC (41). While other studies have demonstrated causal roles of ammonia in promoting the growth of other malignancies (14, 15), our finding of ammonia conferring CSC properties has not been previously reported. Resisting oxidative damage is a hallmark of CSCs that contributes to their long-term self-renewal potential (42) as well as therapy resistance (43). Therefore, anabolic pathways fueled by ammonia-derived nitrogen may have both a direct role in promoting tumor growth and indirect role by helping to maintain low levels of baseline reactive oxygen species.

Our data build on the role of amino acid and nucleotide biosynthesis as drivers of tumorigenesis and directly implicate glutamine-mediated metabolic pathways in contributing to HCC CSC function (44). Targeting CSCs has proven to be a challenge due to some redundancy with other cell populations, but exploiting their distinct metabolic requirements holds potential in the development of new therapeutic applications (42). Several inhibitors of glutamine and related pathways exist that may be explored to address therapy resistance in HCC in subsequent work (45). Given the multifaceted role of glutamine metabolites in biosynthetic pathways, redox balance, cell signaling, and gene expression our data provide a mechanistic understanding of how HCC CSCs may adapt to their microenvironment to promote tumor-propagation.

We demonstrated that SLC4A11 is upregulated in HCC CSCs where it functions as an ammonia importer. While much of the literature has focused on SLC4A11 mutations and their contribution to congenital hereditary endothelial dystrophy (46, 47), recent findings implicate SLC4A11 in HCC by a mechanism that involves ammonia excretion to resist senescence (37). Although the directionality of ammonia transport may be system or cell-type dependent, our conclusions are substantiated by our unbiased tracing of the nitrogen metabolome that demonstrated reduced intracellular nitrogen incorporation in SLC4A11-depleted cells. Our data are consistent in the broader view that SLC4A11 is an important oncogene in HCC. The finding that SLC4A11 confers CSC properties in HCC adds to the understanding of its role in tumorigenesis, warranting further work in HCC and other malignancies.

While this is the largest report to date evaluating clinicopathologic features that correlate with HCC incidence, it remains a retrospective observational study which will require prospective validation of findings. In addition, our work reveals a previously unreported mechanism that involves ammonia-mediated regulation of the nitrogen metabolome that promotes stemness and tumor initiation in HCC. Our clinical data provides correlative evidence that elevated ammonia is associated with higher cancer incidence and poor OS in HCC patients. Targeting ammonia as a therapeutic strategy in HCC has not been explored, and our work suggests the need for clinical trials evaluating whether suppression of ammonia would mitigate HCC incidence or severity in cirrhotic patients.

## Methods

### Sex as a biological variable

Male mice were used in this study given the disproportionate incidence of HCC among males vs females (48, 49). We anticipate that the results of this study are relevant to both genders given its mechanistic basis that applies to HCC initiation in male and female patients.

### Mice

Male 8–12-week-old C57BL/6J and NSG mice were acquired from Jackson laboratories. Mice were fed a standard chow diet ad libitum and housed in a pathogen free, temperature-controlled room with a 12-hour light/dark cycle. Animal experiments were conducted in accordance with the Association for Assessment and Accreditation of Laboratory Animal Care international guidelines and approved by the university committee on the use and care of animals at the University of Michigan.

### Human subjects

This study utilized electronic medical record data from the Veterans Health Administration (VHA) Corporate Data Warehouse (CDW), which includes data for all Veterans receiving care through VHA facilities nationwide. This study was approved by the Veterans Affairs Ann Arbor Research and Development Board. We identified patients diagnosed with cirrhosis from 1999 to 2024 using inpatient/outpatient ICD9/10 codes as well as FIB4 score > 3.25 as previously described (50). The date of cirrhosis diagnosis was defined as the index date in HCC incidence analyses (Cohort 1) and the date of HCC diagnosis was defined as the index date in OS analyses (Cohort 2). Cancer diagnoses and dates of diagnosis were identified by the VA Cancer Registry System. OS was defined as the time from the index date to death from any cause. Date of death was obtained from the VA death registry. Liver function was assessed using ALBI, eCTP, and MELD scores as previously described (24–26). Ammonia levels were obtained from the structured laboratory data. Mean ammonia serum concentration was calculated for patients with greater than one ammonia laboratory value in the system. For HCC incidence analyses, three-year mean ammonia values (one year prior to cirrhosis diagnosis and two years after cirrhosis diagnosis) were used. For OS analyses, two-year mean ammonia values (one year prior to HCC diagnosis and one year after HCC diagnosis) were used. Patients were stratified into low and high ammonia groups based on ≥ 48.5 µM/L which was determined by using the Youden index (51). Propensity score matching was performed via the Toolkit for Weighting and Analysis of Nonequivalent Groups (TWANG) package. The weights were estimated using the covariate balancing propensity score method taking into account age, gender, race, ethnicity, ALBI, eCTP, MELD, Charleson Comorbidity Index, cirrhosis etiology, and HCC screening intensity (by either magnetic resonance imaging or ultrasound). The RADBERT large language model was trained to identify the subset of patients with cirrhosis with negative HCC screens (52). Patients were censored at the date of last known follow-up, defined as the most recent encounter with a VA provider. Patients with ongoing follow-up past April 1, 2024 were administratively censored at that time. Demographics including race, sex, and age were obtained through the Master Patient Index. Analysis was performed with R v4.3.1 (R Core Team, Vienna, Austria) and Python v3.10.4 (Python Software Foundation, Delaware, US).

### Reagents and antibodies

[^14^N]ammonium chloride (Cat. 213330), [^15^N]ammonium chloride (Cat. 299251), [^14^N]ammonium acetate (Cat. A1542), and L-ornithine monohydrochloride (Cat. O6503) were purchased from Sigma. Immunoblotting antibodies were acquired as follows: SLC4A11 (PA5-101889, ThermoFisher Scientific), RFP that cross-reacts with tdTomato (ab124754, Abcam), and actin (3700, Cell Signaling Technologies). CD44 antibody used for flow cytometry was acquired from ThermoFisher Scientific (Cat. 24-0551-82).

### Cell culture

HepG2 cells were provided by Dr. Weiping Zou (University of Michigan) and cultured in Eagle’s Minimum Essential Medium with 10% fetal bovine serum. HUH7 cells were provided by Dr. Susan Uprichard (Loyola University, Chicago) and cultured in Dulbecco’s Modified Eagle Medium with 10% fetal bovine serum. Murine HCC cells (mHCC) were generated from a C57BL/6J mouse bearing HCCs generated by hydrodynamic transfection of Myc, gp53/Cas9, and sleeping beauty transposase (as described below) and were provided by Dr. Viraj Sanghvi (Columbia University) (28). mHCC cells were cultured in Dulbecco’s Modified Eagle Medium with 10% fetal bovine serum. All cell lines were screened for mycoplasma at least bi-weekly and tested negative before use.

### Constructs and Crispr/Cas9 knockout studies

CHOPCHOP (https://chopchop.cbu.uib.no/) was used to design gRNAs for knockout of SLC4A11 in human and murine cell lines. VectorBuilder (https://en.vectorbuilder.com/) was used to clone gRNA sequences into pLV lentiviral vectors with a puromycin resistance cassette. The following gRNA sequences were used: human SLC4A11 #1 5’-AAGGCGATATCCGAGAACA-3’ (VectorBuilder ID: VB231017-1759pac), human SLC4A11 #2 5’-TCGCAGAATGGATACTTCG-3’ (VectorBuilder ID: VB231017-1760qhc), human SLC4A11 #3 5’-GTCCGCAGCACGTTATCCA-3’ (VectorBuilder ID: VB231017-1761uzc), murine SLC4A11 #1 5’-ATTCCAATCCGGTATGACA-3’ (VectorBuilder ID: VB231017-1184qmw), murine SLC4A11 #2 5’-TACTGCACCCCTCGGACAG-3’ (VectorBuilder ID: VB231018-1344xta), murine SLC4A11 #3 5’-GTCCGTGCACACCGGGACC-3’ (VectorBuilder ID: VB231017-1758mtd), and scramble gRNA 5’-TGTAGTTCGACCATTCGTG-3’ (VectorBuilder ID: VB231017-1185cny). Streptococcus pyogenes Cas9-high fidelity variant 1 (SpCas9-HF1) was cloned into a pLV lentiviral vector with a hygromycin resistance cassette (VectorBuilder ID: VB231012-1059dhf) (53). Cells were co-transduced with a gRNA and SpCas9-HF1 and antibiotic selected to generate stable knockout lines. Murine SLC4A11 was connected to tdTomato by a 3X GGS linker at its C-terminus and used for ectopic expression studies (VectorBuilder ID: VB231018-1359kbx). Plasmids used for hydrodynamic transfection (Myc, gp53/Cas9, and sleeping beauty transposase) were provided by Dr. Viraj Sanghvi (Columbia University) and have been previously described (28).

### Hepatosphere assay

Approximately 2,000 cells were plated in ultra-low attachment 6-well plates (Corning, Cat. 3471) in triplicate in advanced Dulbecco’s Modified Eagle Medium/F12 supplemented with 20 ng/mL human recombinant Epidermal Growth Factor (Stem Cell Technologies, Cat. 78136), 20 ng/mL human recombinant Fibroblast Growth Factor (Stem Cell Technologies, Cat. 78134), and 0.2% B27 (ThermoFisher Scientific, Cat. 17504044). Hepatosphere number was quantified when spheres reached at least 100 µM diameter and representative brightfield micrographs were captured using a BioTek BioSpa 8 automated incubator. ImageJ (https://imagej.net/ij/) was used to quantify hepatosphere diameter in brightfield micrographs.

### Real time quantitative PCR

RNA was extracted using an RNA isolation kit (Qiagen, Cat. 74134) and cDNA was produced using high-capacity cDNA reverse transcription kit (ThermoFisher Scientific, Cat. 4368814). SYBR green was used as the master mix (ThermoFisher Scientific, Cat. A25742). Experiments were normalized to glyceraldehyde-3-phosphate dehydrogenase (GAPDH) and performed in triplicate. The following human primer sequences were used: GAPDH: Forward 5’-GGAGCGAGATCCCTCCAAAAT-3’, Reverse 5-GGCTGTTGTCATACTTCTCATGG-3’; SLC4A11: Forward 5’-ATGTCGCAGAATGGATACTTCG-3’, Reverse 5’-AAAAACGGATACTCTCGCCAG-3’; CD44: Forward 5’-CTGCCGCTTTGCAGGTGTA-3’, Reverse 5’-CATTGTGGGCAAGGTGCTATT-3’. The following murine primer sequences were used: GAPDH: Forward 5’-TGGCCTTCCGTGTTCCTAC-3’, Reverse 5’-GAGTTGCTGTTGAAGTCGCA-3’; SLC4A11 Forward 5’-CAGGACTCCGGTGAATACTTCT-3’, Reverse 5’-GATGCTCTCGCCAGACACAA-3’; CD44 Forward 5’-TCGATTTGAATGTAACCTGCCG-3’, Reverse 5’-CAGTCCGGGAGATACTGTAGC-3’.

### ALDH activity assay

Samples were processed for ALDH activity using a colorimetric kit (Sigma, Cat. MAK082). All experiments were performed in triplicate and normalized to the control condition to quantify relative changes in ALDH activity.

### Flow cytometry

Hepatospheres were dissociated into single cells using an enzyme-free dissociation reagent (StemCell Technologies, Cat. 100-0485), processed for CD44 flow cytometry using a Fortessa equipped with four lasers (BD Bioscience), and analyzed using FlowJo software (https://www.flowjo.com/).

### Immunoblotting

Cells were washed in 1x phosphate-buffered saline and lysed using radioimmunoprecipitation assay buffer (ThermoFisher Scientific, Cat. 89900) supplemented with protease (Sigma, Cat. 11836153001) and phosphatase (ThermoFisher Scientific, Cat. A32957) inhibitors. Laemmli 4x sample buffer (BioRad, Cat. 1610747) was added to each sample. The protein lysate was subsequently boiled for 10 minutes at 100°C and separated using SDS– polyacrylamide gel electrophoresis.

### Ammonia quantification

Samples were deproteinated using a methanol-chloroform-water gradient as previously described (14). An ammonia assay kit (Sigma, Cat. MAK310) was used to quantify ammonia concentration in cells or tissues.

### Nitrogen tracing and mass spectrometry

Hepatospheres were generated from HepG2 cells expressing either a scramble gRNA or SLC4A11 gRNA #2. Hepatospheres plates were randomized to control vs ^15^NH_4_Cl treatment (10 mM). After 8 hours of ^15^NH_4_Cl treatment, cells were harvested and pellets were frozen in −80 °C. After conducting 3 independent biological replicates, frozen pellets were processed for HPLC-MS.

For signal normalization, an internal standard (IS) solution containing valine-d_8_, creatinine-d_3_, glutamine-d_5_, phenylalanine-^13^C_6_, and isoleucine-d_10_ (100 μg/mL) was prepared in water/methanol, 1:1 (v/v). Sample extraction was accomplished by adding 40-60 µL IS solution followed by 750-1000 µL water/methanol, 2:8 (v/v). Protein was precipitated by sonication and centrifugation, and samples were transferred to an HPLC autosampler vial and injected directly for LC-MS analysis. Data analysis was performed using Xcalibur Quan Browse software (Thermo; version 4.4.16.14). A custom processing method was created containing all compounds and their isotopologues as previously described (15). A mass accuracy filter of 5 ppm was utilized. Extracted ion chromatograms resulting from the exact m/z values were generated for acidic and basic conditions. Peaks were manually reviewed in both polarities for all samples ran in both mobile phases. Valine-d_8_ was used as the internal standard and the positive-ion data using the acidic LCMS method was used as these generated reliable peak shapes for consistent integration. Data were analyzed using the following formula: Area Ratio Analyte/Internal Standard (x 1,000). This value was used to calculate isotopologue enrichment (%) by quantifying the ratio of the analyte to that of the analyte and its M+0 isotopologue. Respective untreated controls were subtracted from each condition. In some cases, analytes sharing the exact same m/z value were not resolved chromatographically and could not be integrated separately. These analytes are reported in the same rows in the respective figures.

### Metabolite set enrichment analysis

MSEA was performed using MetaboAnalyst 6.0 (https://www.metaboanalyst.ca/). Compounds with ^15^N labeling were analyzed by the KEGG and SMPDB databases and significantly enriched pathways by hypergeometric testing are depicted.

### Limiting dilution analysis

Control and ammonia-treated hepatospheres were dissociated into single cells using an enzyme-free dissociation reagent (StemCell Technologies, Cat. 100-0485), counted, and implanted subcutaneously with *X-vivo* media (Lonza, Cat. 04-380Q) into NSG mice with Matrigel. Control hepatospheres were implanted on the left and ammonia-treated hepatospheres were implanted on the right to assess the effects of each condition in the same mouse. Extreme limiting dilution analysis was used to quantify the tumor initiating cell frequency (35).

### Hydrodynamic and xenograft studies

Hydrodynamic tail vein injections were performed in wild-type male C57BL/6J mice and have been previously described (28). Briefly, a 2 mL plasmid mix of Myc transposon (10 µg), p53 gRNA (10 µg), Cas9 (10 µg), and sleeping beauty transposase (4 µg) was injected into a single mouse. For xenograft experiments, 1 million mHCC cells were subcutaneously implanted in NSG mice with *X-vivo* media (Lonza, Cat. 04-380Q) with Matrigel. Ornithine was delivered by intraperitoneal injection in C57BL/6J mice or subcutaneous implantation in NSG mice at a concentration of 20 mM as previously described (14).

### Ammonium acetate diet

Powdered chow was mixed with 25% ammonium acetate and water using a KitchenAid food mixer as previously described (14). Food pellets were generated and dried in a dehydrator for 72 hours before use.

### Quantification and statistical analysis

Experimental conditions were performed in triplicate and reproducibility of each panel is indicated in the respective figure legend. Representative data is displayed in the figures unless otherwise noted. Mice were randomized to each experimental condition. Details on statistical analyses are described in the figure legends and are reported as the mean ± standard deviation. Statistical significance between two groups was determined by two-tailed *t* test. Statistical significance is described in the figure legends as: * *p* ≤ 0.05, ** *p* ≤ 0.005, *** *p* ≤ 0.0005.

### Study approval

Vertebrate animal studies were carried out under an animal protocol approved by the University of Michigan IACUC.

### Data availability

Reagents that were generated throughout this study are available from the lead contacts with a completed Materials Transfer Agreement.

## Author contributions

A.L.E., M.D.G, and Y.M.S. designed experiments and wrote the manuscript. A.L.E. executed experiments. M.O.E., J.J., Z.W., A.N.P., E.A.H., and A.K.H. provided experimental support. M.G. performed HPLC-MS. T.S.L. provided expertise throughout the entirety of the project and helped write the manuscript H.N.B., V.R.S, T.L.F., G.L.S., E.B.T., A.W.T., N.R., C.P.C., I.D., J.A.M., A.K.B., D.A.E., E.C., J.R.E., K.C.C., T.J.F, D.R.W., M.A.M., D.T.C., and M.S.W. provided feedback on data. All authors read and approved the final manuscript.

## Acknowledgements

The authors thank Dr. Andrew Scott for the helpful discussions.

