## Supplementary Figures 1-5 and Supplementary Table 1 for "SLC4A11 mediates ammonia import and promotes cancer stemness in hepatocellular carcinoma"

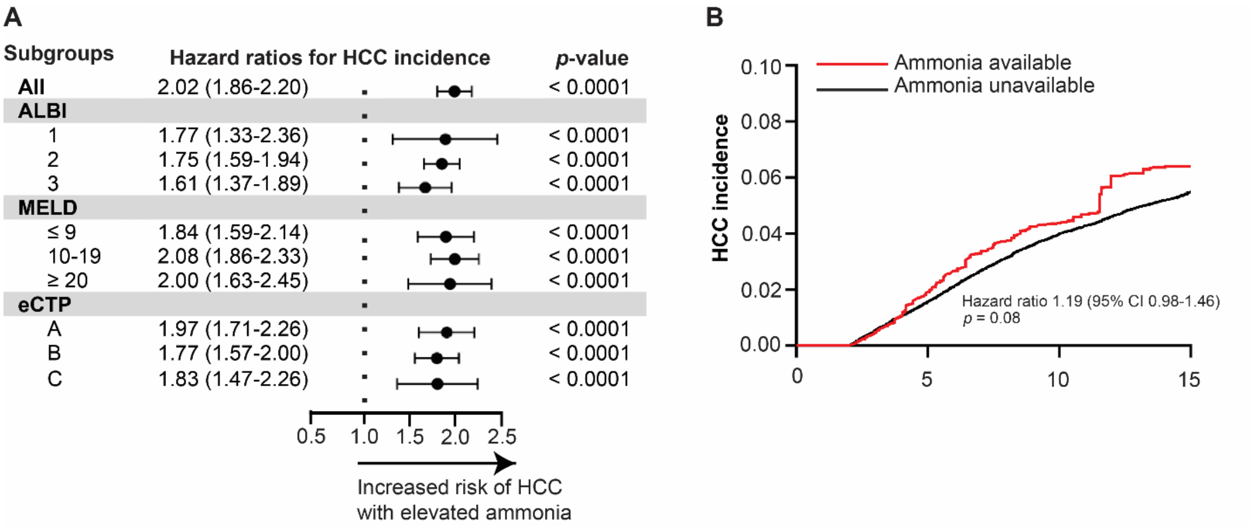


**Supplemental Figure 1.** (**A**) HCC incidence by ALBI grade, MELD score, and eCTP class. (**B**) Two-year landmark analysis of HCC incidence in patients with available and unavailable ammonia using propensity score matching. Hazard ratio log-rank test, *p* values, and 95% confidence intervals indicated.

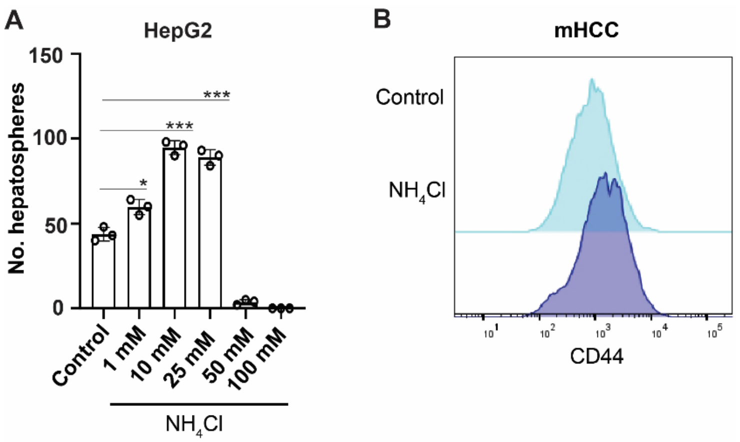


**Supplemental Figure 2.** (**A**) Hepatosphere number of HepG2 cells following treatment with the indicated concentrations of ammonium chloride. Mean ± SD ($n$ = 3). (**B**) Histogram depicting CD44 surface expression in mHCC cells with and without ammonium chloride (10 mM). * $p \leq 0.05$, *** $p \leq 0.0005$ by two-tailed $t$ test.

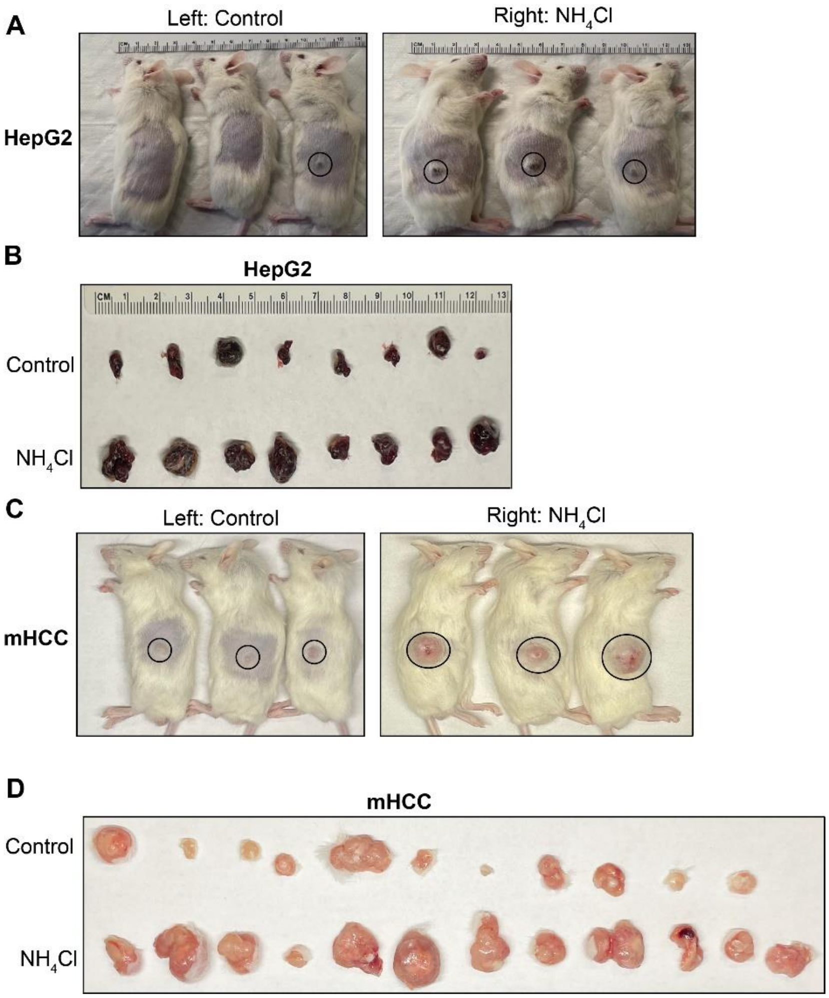


**Supplemental Figure 3.** (**A**) Representative photograph of HepG2 tumor bearing NSG mice with and without pre-treatment with ammonium chloride. (**B**) Representative photo of dissected HepG2 tumors implanted in NSG mice with and without pre-treatment with ammonium chloride. (**C**) Representative photograph of mHCC tumor bearing NSG mice with and without pre-treatment with ammonium chloride. **(D)** Representative photo of dissected mHCC tumors implanted in NSG mice with and without pre-treatment with ammonium chloride.

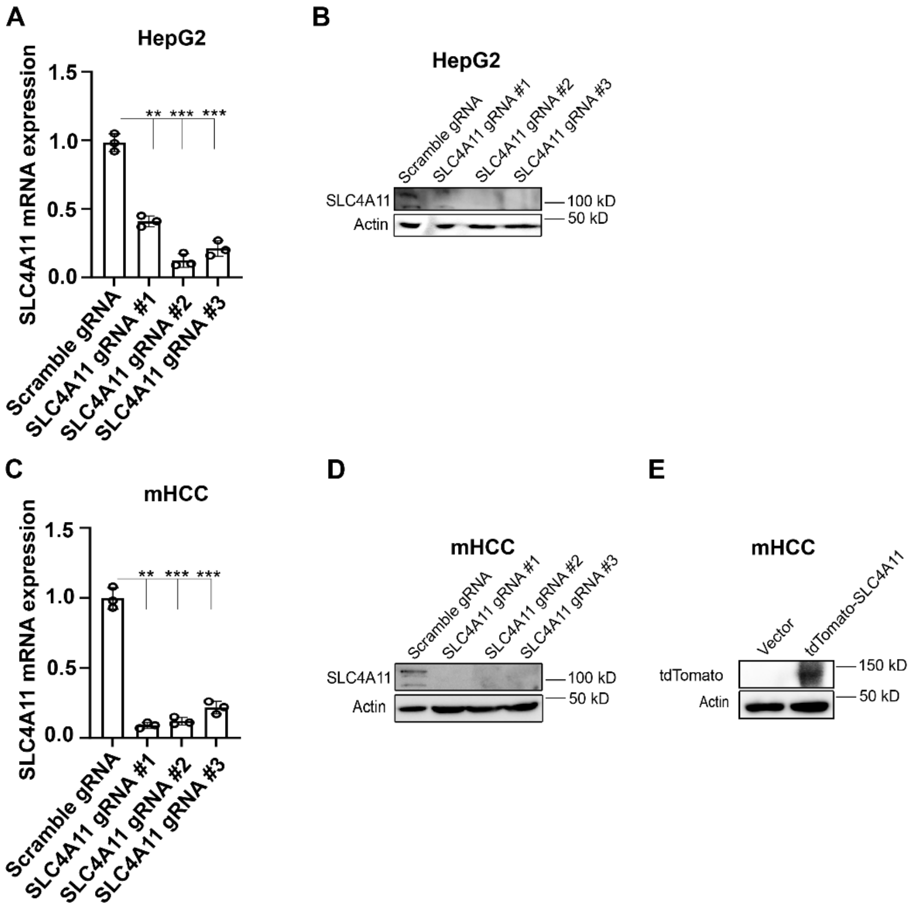


**Supplemental Figure 4.** SLC4A11 was depleted in HepG2 cells by Crispr/Cas9 using 3 independent gRNAs and (**A**) mRNA expression was quantified by qPCR and (**B**) protein abundance was quantified by immunoblotting in control and SLC4A11 KO cells. SLC4A11 was depleted in mHCC cells by Crispr/Cas9 using 3 independent gRNAs and (**C**) mRNA expression was quantified by qPCR and (**D**) protein abundance was quantified by immunoblotting in control and SLC4A11 KO cells. (**E**) tdTomato-tagged SLC4A11 was ectopically expressed in mHCC cells, which was verified by immunoblotting.

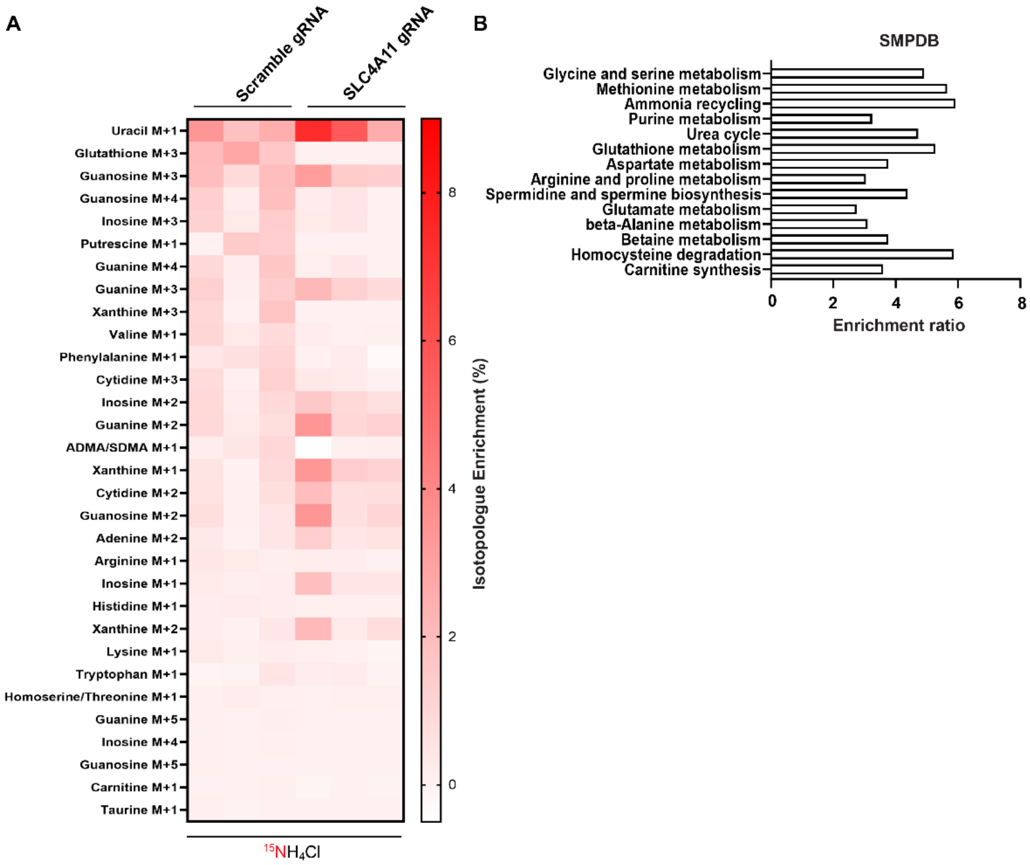


**Supplemental Figure 5.** Isotopologue enrichment in control and SLC4A11 KO HepG2 hepatospheres (background subtracted from $^{15}NH_4Cl$ treated samples, *n* = 3). (**B**) MSEA pathway analysis by SMPDB of 14 pathways determined to be significantly enriched by hypergeometric testing (*n* = 3).

### Supplementary Table 1: Metabolites not detected in $^{15}NH_4Cl$ tracing

| | | | |
|---|---|---|---|
| Acetyl-glycine M+1 | Dihydroorotate M+1 | NAD M+2 | Choline M+1 |
| Adenine M+3 | Dihydroorotate M+2 | NAD M+3 | Niacinamide M+1 |
| Adenine M+4 | Folate M+1 | NAD M+4 | Creatine/hydrolyzed phosphocreatine M+1 |
| Adenine M+5 | Folate M+2 | NAD M+5 | Phosphocholine M+1 |
| Adenylosuccinate M+1 | Folate M+3 | NAD M+6 | NAD M+1 |
| Adenylosuccinate M+2 | Folate M+4 | NAD M+7 | |
| Adenylosuccinate M+3 | Folate M+5 | Niacinamide M+2 | |
| Adenylosuccinate M+4 | Folate M+6 | Nicotinate M+1 | |
| Adenylosuccinate M+5 | Folate M+7 | NMMA M+1 | |
| ADMA/SDMA M+2 | Glucosamine M+1 | NMMA M+2 | |
| ADMA/SDMA M+3 | Glycocholate M+1 | NMMA M+3 | |
| ADMA/SDMA M+4 | GMP M+1 | NMMA M+4 | |
| α-glycerophosphocholine M+1 | GMP M+2 | p-methylhippuric acid/phenylacetylglycine M+1 | |
| AMP M+1 | GMP M+3 | Pantothenate M+1 | |
| AMP M+2 | GMP M+4 | Phosphocreatine dimer/Creatine M+1 | |
| AMP M+3 | GMP M+5 | Phosphocreatine dimer/Creatine M+2 | |
| AMP M+4 | Guanidoacetic acid M+1 | Phosphocreatine dimer/Creatine M+3 | |
| AMP M+5 | Guanidoacetic acid M+2 | Phosphoethanolamine M+1 | |
| Arginine M+2 | Guanidoacetic acid M+3 | Pyridoxine M+1 | |
| Arginine M+3 | Hippurate M+1 | Quinolinate M+1 | |
| Arginine M+4 | Histamine M+1 | Spermidine M+1 | |
| Arginosuccinate M+1 | Histamine M+2 | Spermidine M+2 | |
| Arginosuccinate M+2 | Histamine M+3 | Spermidine M+3 | |
| Arginosuccinate M+3 | Histidine M+2 | Spermine M+1 | |
| Arginosuccinate M+4 | Histidine M+3 | Spermine M+2 | |
| Cadaverine M+1 | Homocysteine M+1 | Spermine M+3 | |
| Cadaverine M+2 | Homocystine M+1 | Spermine M+4 | |
| Carnosine M+1 | Homocystine M+2 | Thymine M+1 | |
| Carnosine M+2 | IMP M+1 | Thymine M+2 | |
| Carnosine M+3 | IMP M+2 | Tryptophan M+2 | |
| Carnosine M+4 | IMP M+3 | UDP-galactose/glucose M+1 | |
| Citrulline M+1 | IMP M+4 | UDP-galactose/glucose M+2 | |
| Citrulline M+2 | Indoleacetic Acid M+1 | UMP M+1 | |
| CMP M+1 | Kynurenic acid M+1 | UMP M+2 | |
| CMP M+2 | Lysine M+2 | Urate M+1 | |
| CMP M+3 | Methionine Sulfoxide M+1 | Urate M+2 | |
| Creatine/hydrolyzed phosphocreatine M+2 | N-acetylleucine M+1 | Urate M+3 | |
| Creatine/hydrolyzed phosphocreatine M+3 | N-acetylputrescine M+1 | Urate M+4 | |
| Creatinine M+1 | N-acetylputrescine M+2 | Xanthine M+4 | |
| Creatinine M+2 | N-acetyltryptophan M+1 | XMP M+1 | |
| Creatinine M+3 | N-acetyltryptophan M+2 | XMP M+2 | |
| Cystathionine M+2 | N6-acetyllysine M+1 | XMP M+3 | |
| Cysteine M+1 | N6-acetyllysine M+2 | XMP M+4 | |
